# Near-Infrared Spectroscopy for metabolite quantification and species identification

**DOI:** 10.1101/277053

**Authors:** WC Aw, JWO Ballard

## Abstract

The aim of the study was to investigate the accuracy of near-infrared spectroscopy (NIRS) in determining triglyceride level and species of wild caught *Drosophila*. NIRS is a remote sensing method that uses the near-infrared region of the electromagnetic spectrum. It detects the absorption of light by molecular bonds and can be used with live insects. We employ the chemometric approach to combine spectra and reference data from a known sample to produce a multivariate calibration model. Once the calibration model was developed, we used an independent set to validate the accuracy of the calibration model. The optimized calibration model for triglyceride quantification yielded an accuracy of 73%. Simultaneously, we used NIRS to discriminate two species of *Drosophila*. Flies from independent sets were correctly classified into *D. melanogaster* and *D. simulans* with accuracy higher than 80%. Finally, we show that the biological interpretations derived from reference data and the NIRS predictions do not differ. These results suggest that NIRS has the potential to be used as a high throughput screening method to assess a live individual insect’s triglyceride level and taxonomic status.

## Introduction

Remote sensing can be defined as any technology that acquires information about an object without making physical contact. Automated machine vision and remote sensing systems are needed and being studied intensively because there are non-invasive, have high-throughput and typically low operating costs (Patel et al., 2012). Benchtop remote sensing acquires data in a controlled environment (e.g., precise lighting, scanning angle, distance between object and probes) and is highly suitable for integration into entomological research. Here, we investigated the potential for benchtop remote sensing, near-infrared spectroscopy (NIRS) to be a non-invasive and high throughput screening method for assessing the triglyceride levels and taxonomic status of two wild-caught *Drosophila* species, *D. melanogaster* and *D. simulans*. *D. melanogaster* and *D. simulans* are globally sympatric sibling species (Capy et al., 2012).

NIRS is a remote sensing technology that detects the “radiometric fingerprint” of a sample by measuring the amount of near-infrared energy absorbed at specific wavelength by biological materials (Manley, 2014). The absorption is influenced by the internal and external chemical composition of the organism and is mainly generated from the stretching and bending of O-H, N-H and C-H functional groups (Williams and Norris, 1987). There are at least five advantages of using NIRS as a remote sensing technology in entomological research. First, it allows simultaneous analysis of multiple components from a single spectrum. Second, the operating cost for NIRS is low as no reagents or sample-specific preparations are needed. Third, NIRS is a high-throughput instrument in which more than 1000 samples can be scanned per day. Fourth, NIRS technology is non-invasive and does not require a highly skilled technician for the operation of the instrument or the analysis of the acquired data after optimisation. Fifth, living organisms can be sampled.

Monitoring targeted metabolites can help understand how an individual performs in response to environmental variables (Nansen and Elliott, 2016). We test the accuracy of NIRS in determining triglyceride levels. Triglycerides constitute the main lipid form, representing ∼90% of the total fat body lipids in insects (Arrese et al., 2006). The content of triglycerides is influenced by several factors, including development stage, nutritional state, sex, and flight activity. Currently, triglycerides in insects are measured by either using commercial assay kits, gas chromatography mass spectroscopy, or liquid chromatography mass spectroscopy (Tennessen et al., 2014). However, all these technologies are costly, invasive, time-consuming and destructive.

Accurate insect species identification is essential for agricultural, ecological, medical, taxonomic, veterinary studies and for biosecurity (Falk et al., 2011; Nansen and Elliott, 2016). Increasingly NIRS is being used by the entomological community and has been shown to accurately identify a range of species including *Anopheles* mosquitoes (Mayagaya et al., 2015), *Zootermopsis* termites (Aldrich et al., 2007), *Calliphoridae* blowflies (Voss et al., 2017), and *Tetramorium* ants (Kinzner et al., 2015). Morphological identification requires taxonomic expertise and is both labor-intensive and time-consuming. A biochemical alternative is to use PCR with direct sequencing or allele specific PCR. Likely, a combined approach will serve as the gold standard for accurate identification of sibling species, however, the processing time and reagent costs often limit their application. As a consequence, field studies of wild caught insects may be based upon a sub sample of all collected individuals, which may not capture the true heterogeneity of the species diversity. Therefore, there is a need for developing effective, low cost and efficient approaches that can be used in the field. Previously, we successfully employed NIRS to determine the species, gender, age and the presence of *Wolbachia* infections in laboratory *Drosophila* (Aw et al., 2012), but the accuracy of identification of wild-caught flies is unknown.

In this study, we employ the chemometric strategy. Chemometric analyses is defined as the development and application of mathematical and statistical methods to extract useful chemical information from sample measurement (Kowalski and Chemistry, 1977). In the chemometric analysis, the best multivariate calibration model is obtained through step-by-step optimization compared to a known reference. This model is then tested with an independent data set and the accuracy determined (Mayagaya et al., 2009; Williams and Norris, 1987).

The goal of this study is to determine the accuracy of NIRS in determining triglyceride levels in two species of *Drosophila* that were captured from nature. We were able to determine triglyceride levels with about 73% accuracy and species with greater than 80% accuracy. Combined, these results suggest that NIRS has the potential to be used as a high throughput screening method to assess a live individual insect’s triglyceride levels and species status.

## Materials and Methods

The analytical strategy followed in this study is summarized in Fig. 1.

**Fig. 1.**
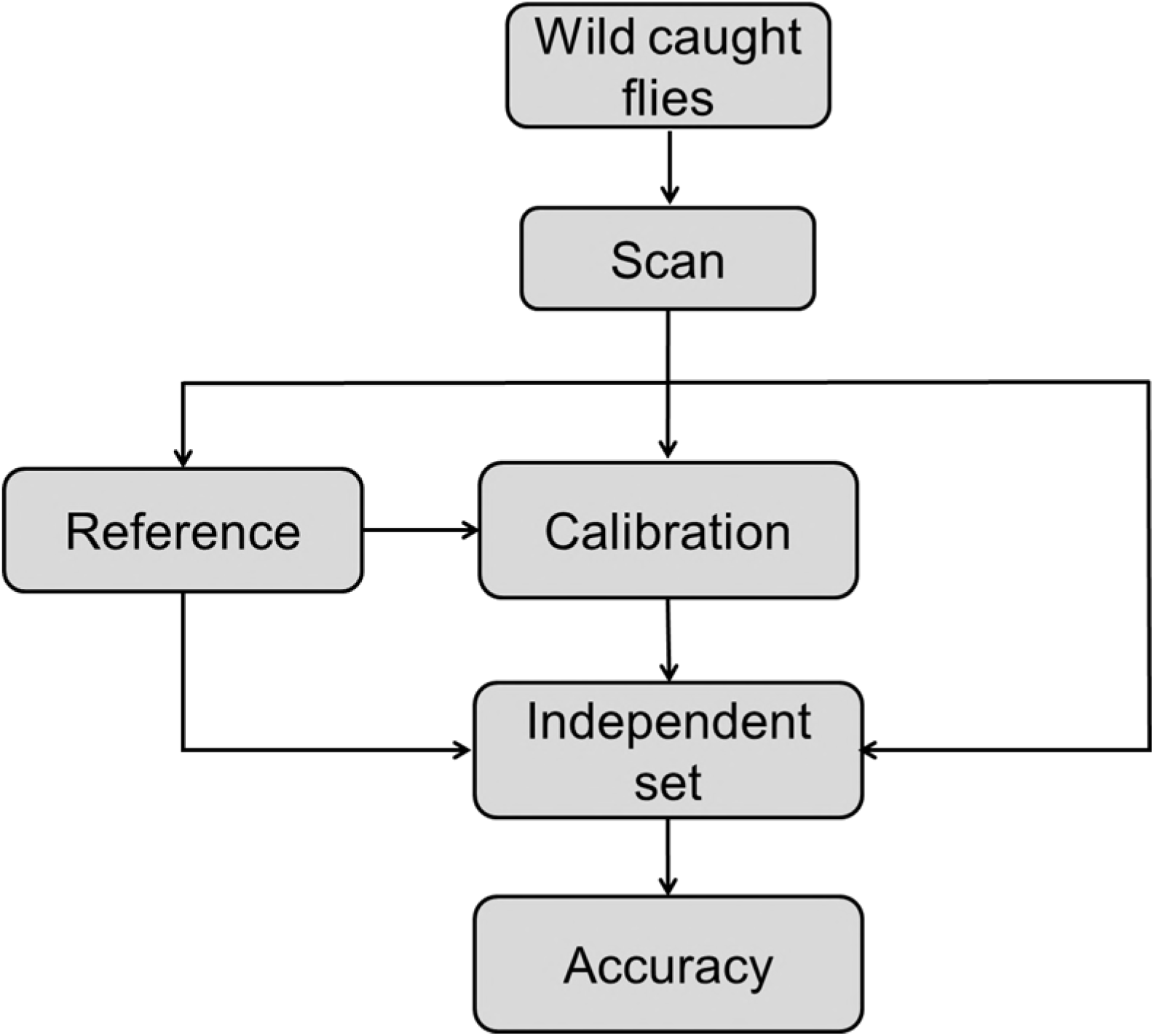
Chemometric analytical strategy for triglyceride quantification and species identification. A total of 280 wild caught flies were included in the study. Of these, 159 were used to construct the calibration models and 121 used to construct the independent sets. After scanning, wild caught flies were used to develop reference data for triglyceride quantification using Sigma triglyceride assay kit or species identification with allele specific PCR of a mtDNA fragment (n= 112 and n= 168 for triglyceride quantification and species identification, respectively). Both methods were destructive and therefore samples used for triglyceride quantification could not be used on species identification. Multivariate calibration models were generated by associating scanned data with reference data (Total 159 flies; n= 65 and n= 94 for calibration model of triglyceride quantification and species identification, respectively). Independent sets build with data not included in the calibration models (Total 121 flies; n= 47 and n= 74 for independent set of triglyceride quantification and species identification, respectively) were used to estimate the accuracy of the calibration model.

### Wild caught flies

Flies used to develop calibration models and validation sets were handled and processed the same way to increase accuracy (Li et al., 2017). Specifically, wild caught *Drosophila* flies were collected in Rosebery, NSW, Australia and scanned with NIRS within three hours of collection. Flies were anesthetized with humidified CO_2_ for 30 min immediately before the scan was performed. After NIRS scanning, flies were frozen in liquid nitrogen and then transferred into a -80° C freezer.

### Scan

The scanning system set-up follows Mayagaya et al. (2009). About 25 flies were placed on a spectralon plate (ASD Inc., Boulder, Colorado, USA) and each fly was individually scanned. The flies were placed 2 mm below a 3 mm diameter bifurcated fiber-optic reflectance probe that contained 33 illumination fibers and 4 collection fibers. The probe was focused dorsally on the head and thorax and the spectra were collected with a portable LabSpec 5000 spectrometer (350-2500 nm; ASD Inc., Boulder, Colorado, USA) using RS^3^ Spectra Acquisition Software 6.0.10 (ASD Inc., Boulder, Colorado, USA). The raw channel data sampling rate of 1.4 nm in the visible and near-infrared region (350 to 1000 nm) and 2.2 nm in the short wavelength infrared region (1001 to 2500 nm) are interpolated to 1 nm intervals across the full spectrometer range from 350 nm to 2500 nm. The nominal spectral resolution varies with the spectrometer region. The visible and near-infrared region has a spectral resolution of 3 nm at 700 nm and the short wavelength infrared region has a spectral resolution of 10 nm at 1400 nm and 2100 nm. An average of 50 spectra was collected from each sample and stored as an average spectrum. All spectra were converted into SPC format by the Asd to Spc convertor version 6 (ASD. Inc.). The spectra were transformed into log 1/R and mean centered before analysis.

### Reference

The reference data for triglyceride level and species status were independently determined. These reference data were fed into the development of the chemometric calibration model and independent data set. In the calculation of the accuracy of NIRS it is assumed that the reference data was obtained without error.

Reference triglyceride levels were determined using the Abcam fluorometric kit (AB65336) following manufacturer’s instructions. Briefly, triglycerides are hydrolyzed to free fatty acids and glycerol. The glycerol reacts with the triglyceride enzyme mix to form an intermediate product, which in turn reacts with the PicoProbe and developer to generate fluorescence that can be detected at Ex/ Em= 537/ 587 nm. Experimental samples were prepared by grinding each adult fly in 100 µl of 5% NP-40/ ddH_2_O. Samples were slowly heated to 85°C for 5 mins and then cooled down to room temperature. The heating and cooling process was repeated twice and samples were centrifuged for 2 min at 4000 g to remove insoluble materials. Triglyceride level was performed in a 384 well microplate, with each well containing 25 µl of samples in 25 µl of working buffer (consists of 23.8 µl of triglyceride assay buffer, 0.2 µl of triglyceride probe and 1 µl of triglyceride enzyme mix). All measures were performed at 23 °C. Triglyceride level was expressed as nmol/ well.

Reference species data was determined with allele specific PCR and the robustness of this protocol was corroborated using laboratory flies of known taxonomic status. Allele-specific PCR primers were designed by downloading and aligning 42 *cytochrome c oxidase I* (*cox I*) sequences from GenBank (15 *D. melanogaster* and 27 *D. simulans*). *Cox I* is a mitochondrial DNA encoded gene which is recognized as an extremely useful DNA barcode, capable of accurate species identification in a broad range of animals (Hebert et al., 2003). *D*. *simulans* was identified by amplifying a 784 bp region of *cox I* gene using primers 1856F (5’- TATCTGCTGGAATTGCCCAC-3’) and 2642R (5’-GCTATAATAGCAAATACAGCTCC-3’), while *D*. *melanogaster* was identified by amplifying a 600 bp region of *cox I* gene using primers 2041F (5’- GCTTTATTATTATTATTATCACTT-3’) and 2642R (5’-GCTATAATAGCAAATACAGCTCC-3’). DNA was extracted from flies using a Gentra Puregene® Cell kit (Gentra Sytem Inc., Minneapolis, MN, USA). Each 10 µl reaction contained 2 µl of Crimson^TM^ buffer (NEB, New England Biolabs), 2.56 µl of 25mM MgCl _2_, 0.4 µl of 10mM forward and reverse primer, 0.08 µl of 25mM dNTP, 0.05 µl of Taq polymerase, 2.51 µl of H _2_O and 10 ng of DNA. The PCR cycling program involved four separate phases. Phase 1 was the initial denaturation which was 94 °C for 2 min. Second, the 5 cycle touchdown phase (denaturation: 94 °C for 10 s, annealing: 64 °C for 15 s with the temperature gradually reducing 1°C per cycle until it reached 59 °C, and extension: 1 min at 72 °C). Third, the 20 cycle phase (denaturation 94 °C for 10 s, annealing 59 °C for 15 s and 1 min at 72 °C). Fourth, a final 72 °C extension step for 6 min.

### Calibration

The calibration models were constructed with partial least square (PLS) regression leave-one-out cross-validation method using GRAMS IQ version 9.1 (Thermo Fisher Scientific, Salem, NH)(Williams and Norris, 1987). This technique is appropriate for small sample sizes. Cross-validation is carried out by dividing the population of samples into equal “blocks” and eliminating samples one block at a time. As a consequence, all samples are used in the development of the calibration equation. A regression coefficient plot was used to identify the important peaks and the noise region in the model. This plot shows that noise detrimental to the model increases outside 500-2200 nm. As a consequence, the region outside 500-2200 nm was excluded from analyses. All spectra were smoothed using the Savitzky-Golay first derivative method (Savitzky and Golay, 1964) and outliers identified by Mahalanobis distance (Mahalanobis, 1936). There were less than 5% outliers in all models.

For triglyceride quantification, a calibration model was generated using 65 wild-collected flies. In the model, the spectra were assigned with the reference value obtained from the fluorometric assay. All spectra were processed using the Savitzky-Golay first derivative with 35 point smoothing function.

For species identification, three calibration models with cross-validation diagnostic methodology were developed. The calibration models were developed using either mixture of male and female flies (94 flies), male flies (50 flies) or female flies (44 flies). In the calibration model, the spectra of *D. melanogaster* flies were assigned a value of 1, and *D. simulans* were assigned a value of 2. The value of 1.5 was considered as the cut-off point for species identification. Flies with predicted value less than 1.5 were classified as *D. melanogaster*, whereas those with a predicted value equal or greater than 1.5 were classified as *D. simulans*. All spectra were processed using the Savitzky-Golay first derivative with 5 point smoothing function.

### Independent set

We used a training data set to construct a calibration model that relates the multivariate response (spectrum) to the concentration of the analyte of interest. A disadvantage of this approaches that search through training data for empirical relationships tend to overfit the data. In order to avoid overfitting, we used the independent set built with data not included in the calibration models to validate each calibration model. If a model fit to the training set also fits the test set well, minimal overfitting has taken place (Subramanian and Simon, 2013). The independent set was analyzed using GRAMS IQ Predict version 9.1 (Thermo Galactic, Salem, NH).

For triglyceride quantification, the independent set was generated using 47 wild-collected flies. The predicted value for triglyceride level was then determined using the calibration model and compared with that determined by the fluorometric assay.

For species identification, three independent sets were used to estimate the accuracy of each calibration model. The independent sets were developed using either a mixture of male and female flies (74 flies), male flies (44 flies) or female flies (30 flies). Spectra with species identified using allele specific PCR were then compared with that predicted from calibration model.

### Accuracy

Accuracy represents the combination of the sum of the trueness (systematic error) and precision (random error) (Baratloo et al., 2015). Here, the best multivariate calibration model was chosen based on the highest accuracy of prediction of the independent set.

Triglyceride levels determined from the fluorescent kit were continuous and the accuracy of the triglyceride level quantification was determined by measuring the root-mean-square error of calibration, coefficient (*R*^*2*^) and root-mean square error of prediction. Root-mean-square error of calibration and *R*^*2*^ were used to measure goodness of fit between the reference data and the calibration model. The root-mean square error of prediction, computed from the independent set, was used to measure the differences between NIR reflectance and the reference value. The closer the predicted scan result is to the actual or known result the lower the root-mean square error of prediction value. A good model should have lower root-mean-square error of calibration, lower root-mean square error of prediction and higher *R*^*2*^. To enable comparison with the species identification results we focus upon *R*^*2*^ as a measure of accuracy.

Species identification using allele specific PCR was discrete (1 for *D. melanogaster* and 2 for *D. simulans*). The accuracy was calculated by comparing the allelic specific PCR result with the scan result. The closer the predicted result is to the allelic specific PCR result the greater the accuracy.

### Statistical anlayses

To verify whether the biological interpretations derived from the reference method and NIRS method is in agreement with each other, we analyses the data obtained from these two methods by ANOVA and Student’s t-tests using JMP 12 Software (© 2015 SAS Institute, Cary, NC, USA). The main effects were species and sex.

### Result and Discussion

One challenge facing many entomological studies is the accurate and cost effective linking of physiological features and species status. In response to this challenge we aimed to determine the accuracy of NIRS by determining triglyceride levels and taxonomic status of wild-caught *Drosophila*.

### NIRS triglyceride quantification

In nature, fat reserves are the most important reserve used by insects to meet their energy demand during diapause (Hahn and Denlinger, 2007) and influences starvation survival (Ballard et al., 2008). Here, reference triglyceride concentrations of the wild caught flies were determined using a commercial fluorometric kit (Fig. S1) and ranged between 0.312-1.526 nmol/ fly.

A regression coefficient plot was used to display the component of interest of the property being investigated. The regression coefficient plot for triglyceride quantification showed peaks in the regions around 1140 nm, 1370 nm, 1820 nm and 1900 nm (Fig. 2) which were consistent with the absorptions of functional groups associated with glycerol and fatty acids. These functional groups include methyl group (-CH_3_), methylene group (-CH_2_), alkene group (C=C), and ester group (COOC). The peaks at 1140nm and 1370nm are characteristic of the 2^nd^ overtone and the combination of C-H stretching. Peak on1820nm shows the 1^st^ overtone of C-H stretching, whereas peak on 1900nm corresponds to the absorption of COOC functional groups (Williams and Norris, 1987).

**Fig. 2.**
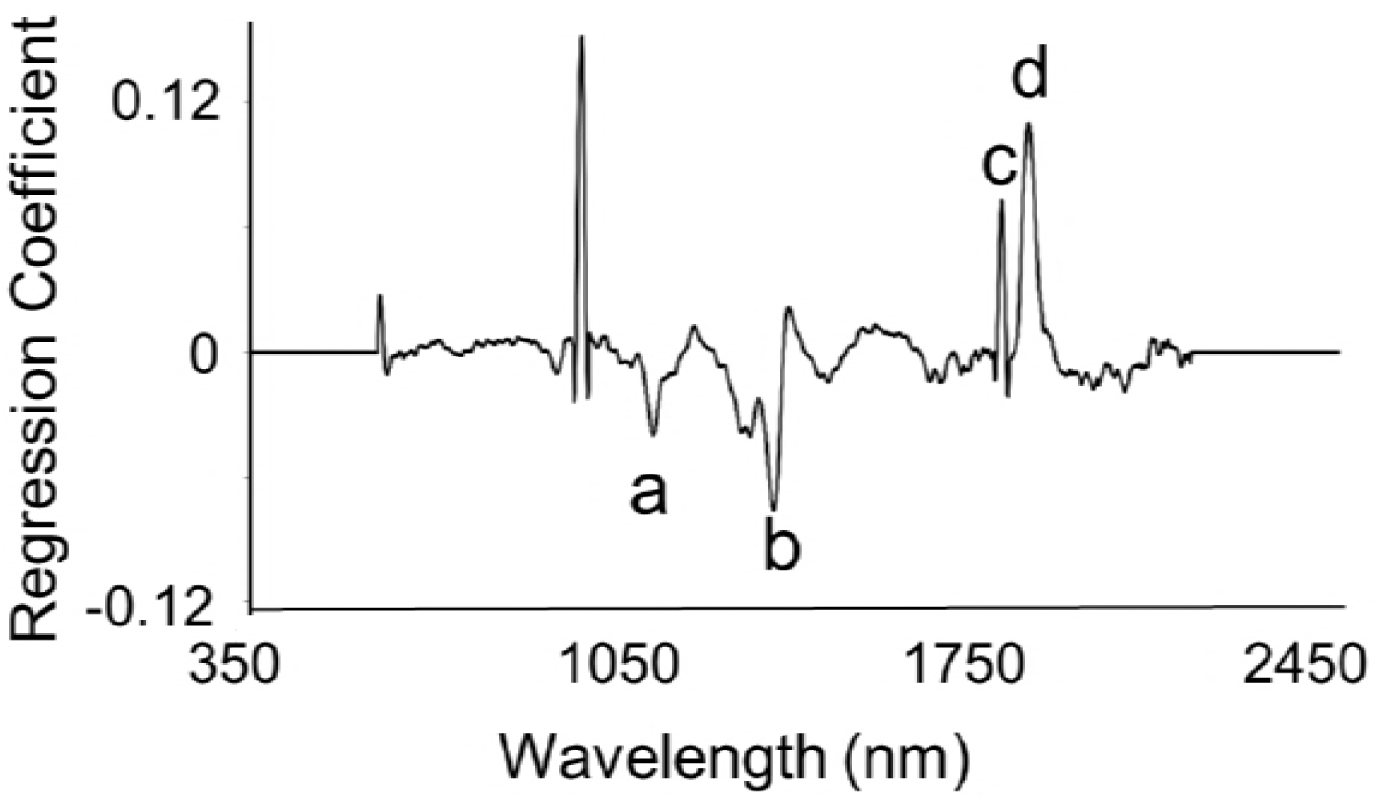
Regression coefficient plot for triglyceride quantification. (a) 1140 nm, (b) 1370 nm,(c) 1820 nm and (d) 1900 nm.

The optimized calibration model for triglyceride quantification yields an accuracy of 73% (Fig. 3), root-mean square error of calibration of 0.19 and root-mean square error of prediction value of 0.26. In contrast to our results, Neves et al. (2012) developed a NIRS calibration model with 96% accuracy for triglyceride quantification of animal plasma. Plausibly, NIRS scanning of insect hemolymph fluid may improve the accuracy of the calibration model.

**Fig. 3.**
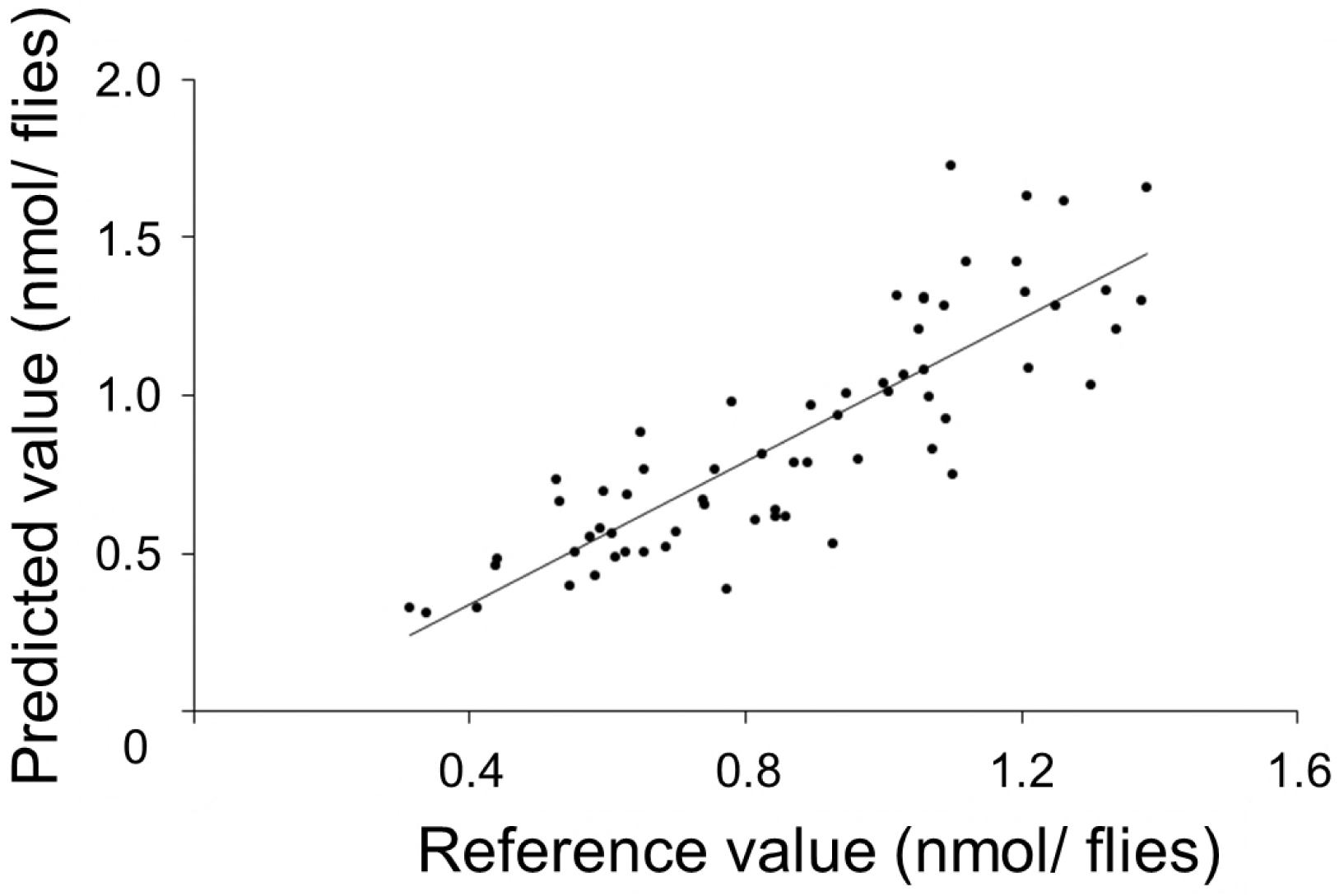
Relationship between the reference fluorometric kit and the NIRS predicted triglyceride values. The calibration model has an 73% accuracy of prediction.

### NIRS species identification

High morphological similarity between sibling species can make identification difficult. Male *D. melanogaster* can be differentiated from male *D. simulans* by the shape of the genital arch genitalia, but females are difficult to identify. In this study, we categorized the wild caught flies as either *D. melanogaster* or *D. simulans* using allele-specific PCR (Fig. S2).

The regression coefficient plot show peaks in the regions around 1450 nm, 1720 nm, 1820 nm and 1900 nm (Fig. 4). Notably, the peak around 1450 nm was observed in our previous study on species identification of laboratory *Drosophila* (Aw et al., 2012). The peak at 1450 nm is characteristics of the 1^st^ overtone and the combination of C-H stretching and has been shown to increase with the rise in moisture content of the sample (Yang et al., 2013). Peaks at 1820 nm and 1900 nm are observed in the regression coefficient plot for triglyceride quantification (Fig. 2). This indicated that lipid’s may also play an important role in species discrimination.

**Fig. 4.**
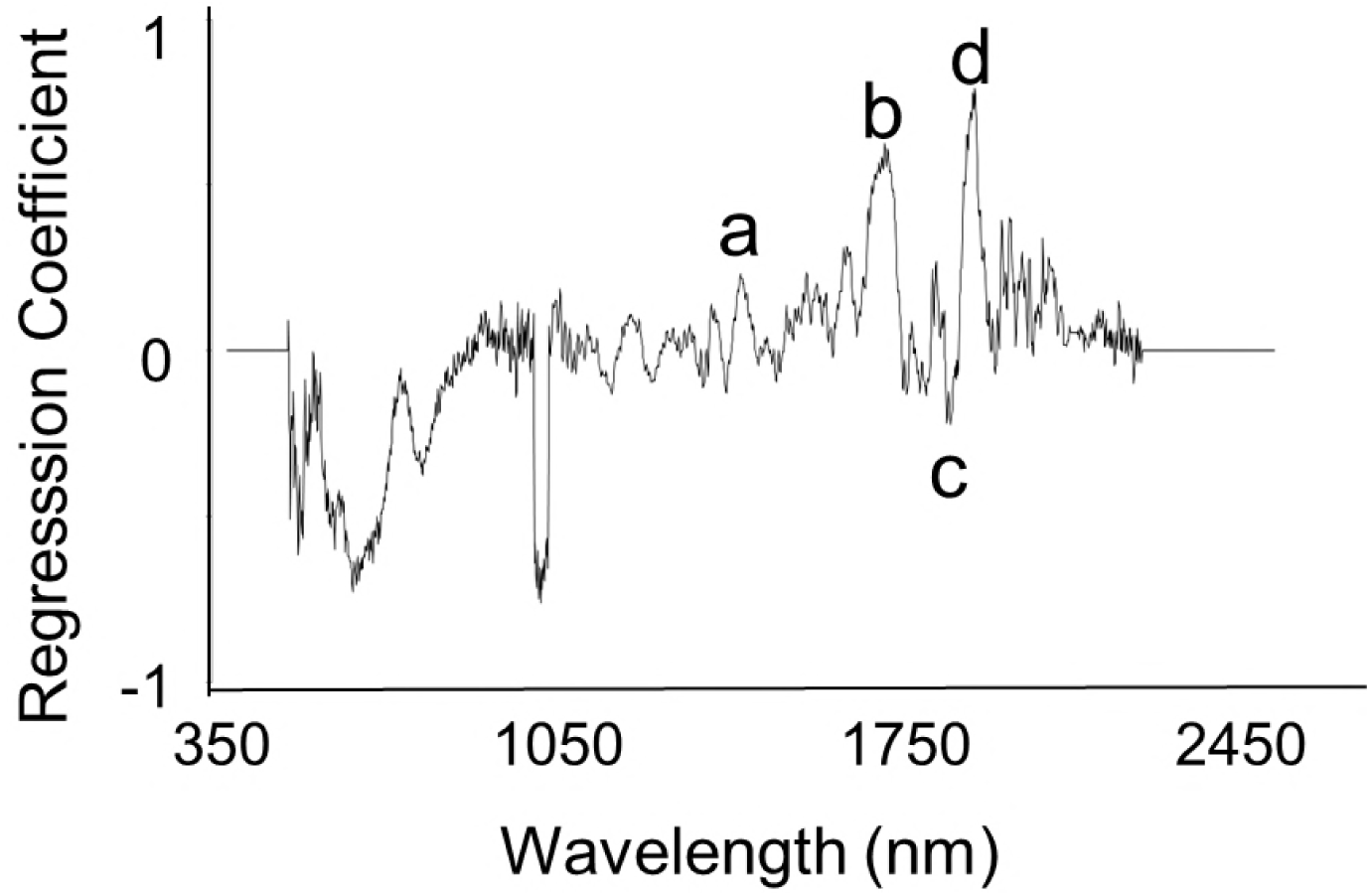
Regression coefficient plot for classifying *D. melanogaster* and *D. simulans.* (a) 1450 nm, (b) 1720 nm, (c) 1820 nm and (d) 1900 nm.

Three calibration models were developed using the mixture of male and female flies (Model 1), male flies (Model 2) and female flies (Model 3). Male and female flies were correctly classified as *D. melanogaster* and *D. simulans* with 80% (n= 74) accuracy, respectively, using calibration Model 1. In calibration Model 2, male flies were correctly classified as *D. melanogaster* and *D. simulans* with 86.4% (n= 22) and 90.9% (n= 22) accuracy, respectively. In calibration Model 3, female flies were correctly classified as *D. melanogaster* and *D. simulans* with 80% (n= 15) and 86.7% (n= 15) accuracy, respectively (Fig. 5).

**Fig. 5.**
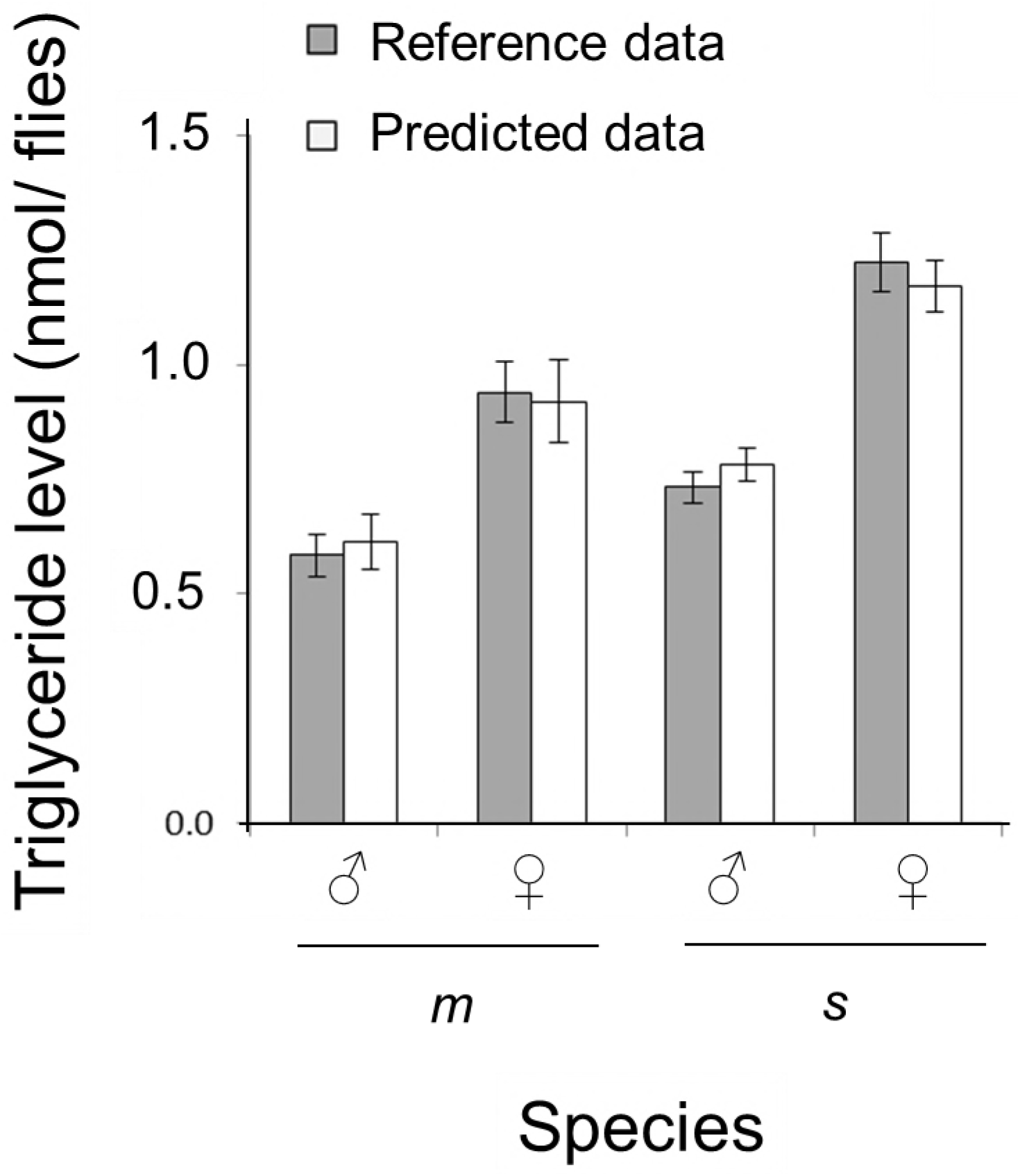
Reference data (grey box) and the NIRS predicted data (white box) for males(**♂**) and females (♀) *D. melanogaster* (*m*) and *D. simulans* (*s*).

**Fig. 5.** NIRS species identification of *D. melanogaster* (*m*) and *D. simulans* (*s*) using three calibration models. Dotted line indicated cut-off point for delineating species. Calibration Model 1 includes all flies. Calibration Model 2 includes male flies. Calibration Model 3 includes females.

The optimized calibration model for species identification of wild caught *Drosophila* (Fig. 5) and of laboratory reared *Drosophila* (Aw et al., 2012) had greater than 80% accuracy of prediction. Similarly, the NIRS calibration model for two field collected mosquito species also had an accuracy of approximately 80% (Mayagaya et al., 2009). In comparison, the accuracy of identifying four *Tetramorium* ant species was lower (13.3% - 66.7%) (Kinzner et al. (2015). Future studies should consider how the accuracy of NIRS species identification could be improved.

### Comparison of observed and predicted data

Finally, we wished to see if the biological interpretations derived from the reference data (flourometric and allele specific PCR) and the results predicted from NIRS would differ. The data suggests that females have higher triglyceride levels than males and *D. melanogaster* has lower levels than *D. simulans* (Fig. 5). ANOVA of the reference data shows significant effects of species (F_1, 61_= 11.65, p = 0.001) and sex (F_1, 61_= 42.07, p < 0.001), but no significant species by sex interaction (F_1, 61_= 1.24, p = 0.270). For the NIRS method, ANOVA also reported significant effects of species (F_1, 61_= 7.52, p = 0.008) and sex (F_1, 61_= 34.55, p < 0.001) but no significant sex by species interaction (F_1, 61_= 0.02, p = 0.877).

Student’s t tests showed no significant differences between the methodologies at the 95% confidence level for *D. melanogaster* (t_30_ = 0.369, p= 0.714; t_10_ = 0.429, p= 0.677, respectively for males and females) or *D. simulans* (t_44_ = 0.354, p= 0.937; t_38_ = 0.560, p= 0.604, respectively for males and females).

Caution must be exercised in over interpreting the biological implications of the results because multiple variables can affect triglyceride content in field collected flies. However, the data predict that females of both species should be more resistant to starvation than males and *D.simulans* should be more resistant to starvation than *D. Melanogaster.* The former postulation is supported by Ballard et al. (2008) and Robinson et al. (2000), who showed female *Drosophila* are more starvation resistant than males. However, our prediction conflicts with Behrman et al. (2015) who showed *D. melanogaster* are more starvation resistant than *D. simulans*. In addition to the ecological and behavioural differences that may effect triglyceride levels between species, it is also possible that starvation resistance is highly genotype specific.

## Conclusion

The current methods for metabolite quantification and sibling species identification can be difficult and laborious. To overcome these difficulties, we tested the potential use of NIRS for triglyceride quantification and species identification. The major advantages of NIRS technique for entomologists include the cost saving after initial purchase of the instrument, nondestructive sampling and the potential for high-throughput analysis. The major limitation is that the methodology was not 100% accurate. Our study demonstrated NIRS can quantify triglyceride and identify wild-caught *Drosophila* with greater than 73% accuracy. In cases where high accuracy is required, NIRS may be able to provide an initial screening of the data as the specimens are not damaged.

Ongoing goals are to increase the accuracy and usage of NIRS. NIRS may potentially include certain artefact peaks in the calibration models and lead to inaccuracy of prediction. Instrument noise is one of the major sources of inaccuracy that can be reduced by proper calibration of the instrument. The other artefacts could be as a result of non-controlled conditions (e.g., lighting, scanning angle, distance between object and probes).

Additional challenges include linking additional metabolites with NIRS spectral patterns and simultaneously identifying more than two species. Kinzner et al. (2015) demonstrate that four species of ant (*Tetramorium*) could be classified by NIRS using one-vs-all strategy with an accuracy of 13 – 67% (Rifkin and Klautau, 2004). A major limitation of Kinzner et al. (2015) model is that the accuracy of prediction is low. Finally, our study provides evidence that NIRS is a promising method for monitoring of insect’s metabolite level and taxonomic status. In addition to insect’s studies, we believe that this technology can be used in medical field for diagnosis of diseases in case where it is known that specific metabolites differ.

## Acknowledgements

We thank the Ballard group for comments, Gary Fager and Paul Martin (ASD Inc.) for technical support. Funding was provided by the Australian Research Grant LE110100134 toJ.W.O. Ballard.

---

**Fig. S1.**
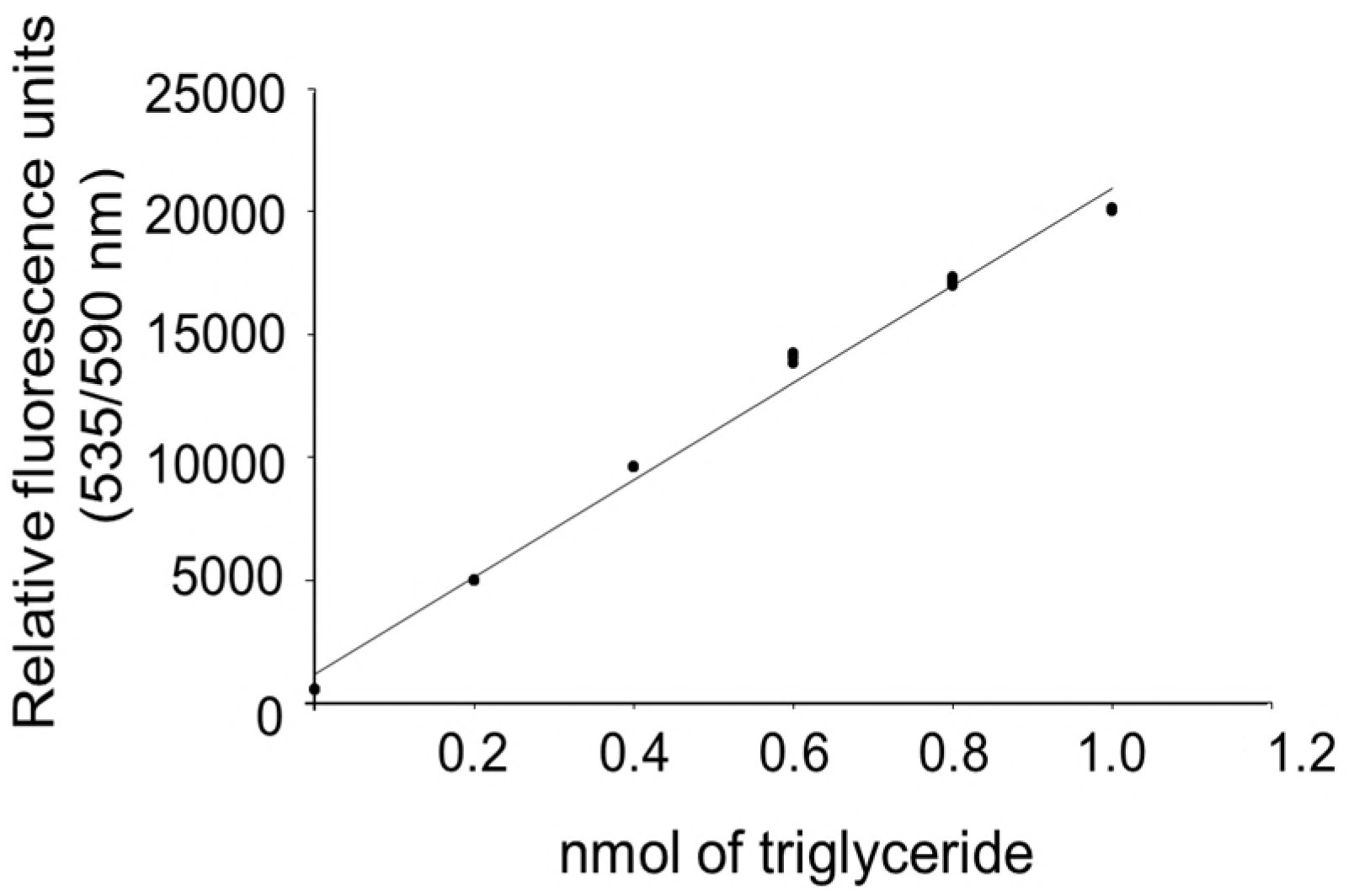
Flurometric triglyceride standard curve. The standard curve has a 99% accuracy of prediction.

**Fig. S2.**
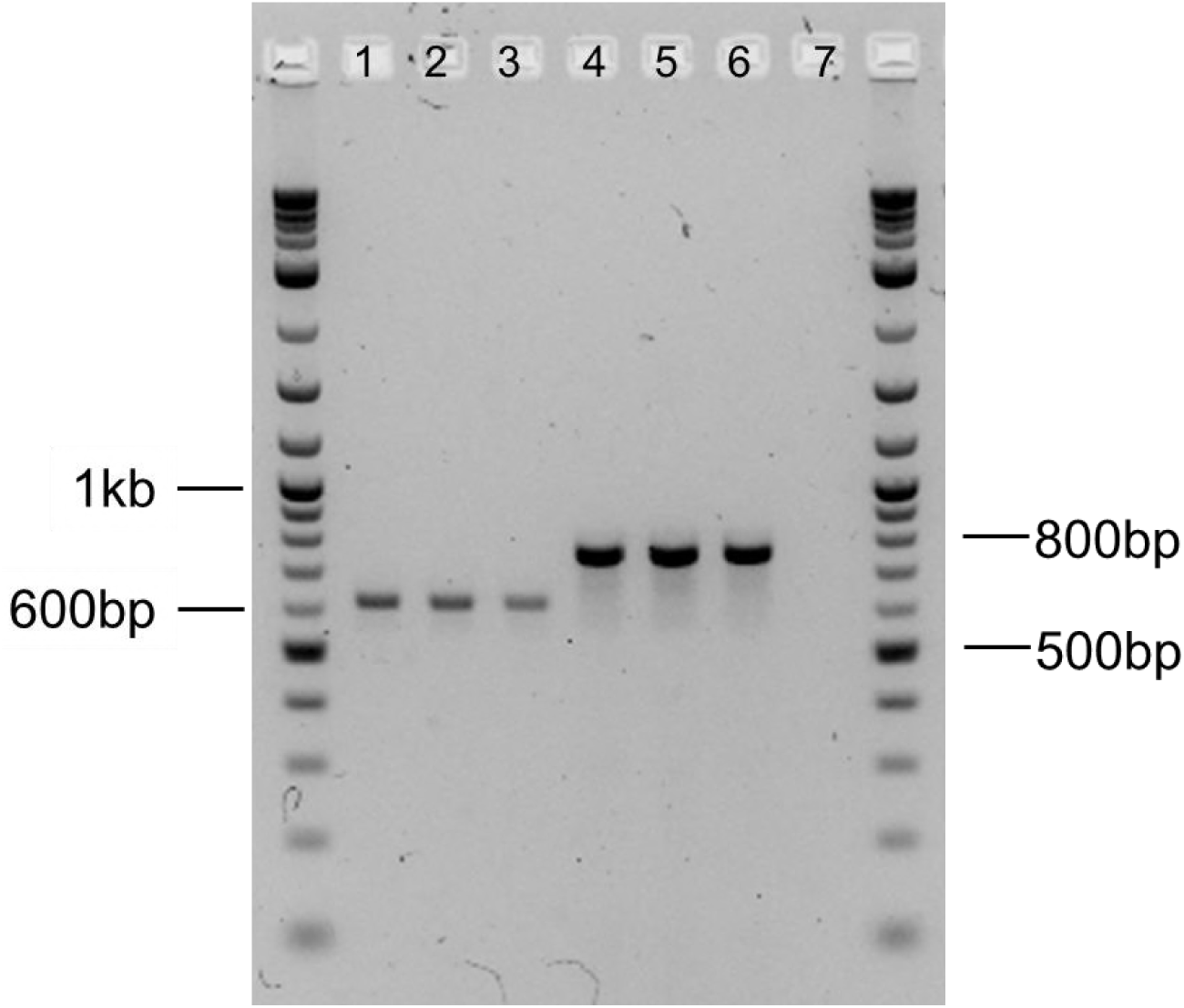
Multiplex allele-specific PCR for species determination. Primer for allele-specific PCR was designed to amplify 600 bp of *D. melanogaster* mtDNA *cox I* gene (well 1, 2 & 3) and 784 bp of *D. simulans* mtDNA *cox I* gene (well 4, 5 & 6). The negative control was located in well 7. A 100 bp ladder is located on both sides of the gel.

